# A neural algorithm for a fundamental computing problem

**DOI:** 10.1101/180471

**Authors:** Sanjoy Dasgupta, Charles F. Stevens, Saket Navlakha

## Abstract

Similarity search, such as identifying similar images in a database or similar documents on the Web, is a fundamental computing problem faced by many large-scale information retrieval systems. We discovered that the fly’s olfactory circuit solves this problem using a novel variant of a traditional computer science algorithm (called *locality-sensitive hashing*). The fly’s circuit assigns similar neural activity patterns to similar input stimuli (odors), so that behaviors learned from one odor can be applied when a similar odor is experienced. The fly’s algorithm, however, uses three new computational ingredients that depart from traditional approaches. We show that these ingredients can be translated to improve the performance of similarity search compared to traditional algorithms when evaluated on several benchmark datasets. Overall, this perspective helps illuminate the logic supporting an important sensory function (olfaction), and it provides a conceptually new algorithm for solving a fundamental computational problem.

---

An essential task of many neural circuits is to assign neural activity patterns to input stimuli, so that different inputs can be uniquely identified (*1, 2*). Here, we study the circuit used by the fruit fly olfactory system to process odors and uncover new computational strategies for solving a fundamental machine learning problem: approximate similarity (or nearest-neighbors) search.

The fly olfactory circuit assigns each odor a “tag”, corresponding to a set of neurons that fire when that odor is presented (*3*). This tag is critical for learning behavioral responses to different odors (*4*). For example, if a reward (e.g., sugar water) or a punishment (e.g., electric shock) is associated with an odor, that odor becomes attractive (a fly will approach the odor) or repulsive (a fly will avoid the odor), respectively. The tags assigned to odors are known to be sparse — only a small fraction of the neurons that receive odor information respond to each odor (*5, 6*) — and non-overlapping — tags for two randomly selected odors share few, if any, active neurons, so that different odors can be easily distinguished (*3*).

The tag for an odor is computed using a three step procedure (Figure 1A). The first step involves feed-forward connections from odorant receptor neurons (ORNs) in the fly’s nose to projection neurons (PNs) in structures called glomeruli. There are 50 ORN types, each with a different sensitivity and selectivity for different odors. Thus, each input odor has a location in a 50-dimensional space determined by the 50 ORN firing rates. For each odor, the distribution of ORN firing rates across the 50 ORN types is exponential with a mean that depends on the concentration of the odor (*7, 8*). For the PNs, this concentration-dependence is removed (*8–12*); i.e., the distribution of firing rates across the 50 PN types is exponential with close to the same mean for all odors and all odor concentrations (*3*). Thus, the first step in the circuit essentially “centers the mean” — a standard pre-processing step in many computational pipelines — using a technique called divisive normalization (*13*). This step is important so the fly does not mix up odor intensity with odor type.

**Figure 1:**
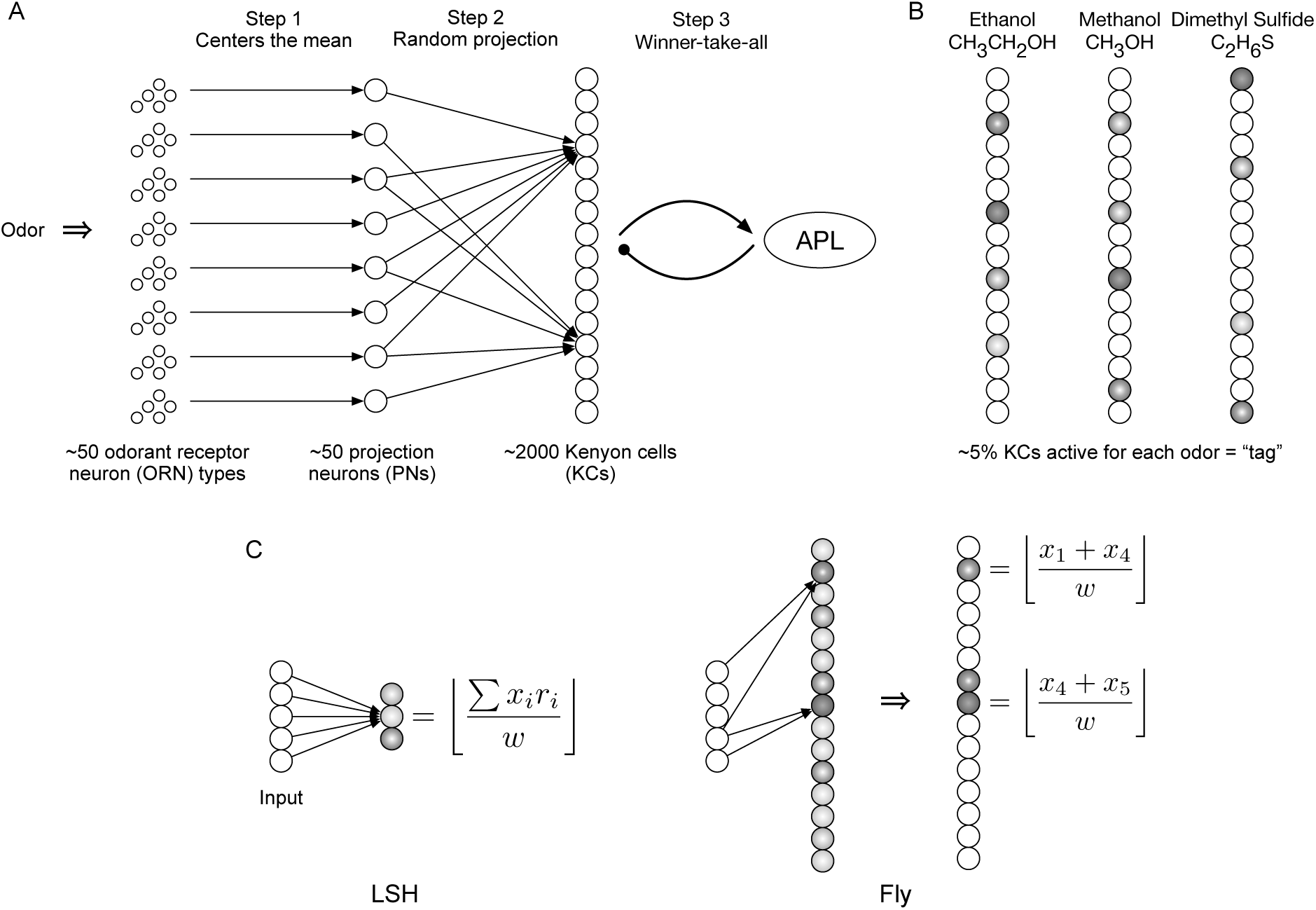
Mapping between the fly olfactory circuit and locality-sensitive hashing (LSH). A) *Schematic of the fly olfactory circuit.* In Step 1, 50 ORNs in the fly’s nose send axons to 50 PNs in the glomeruli; as a result of this projection, each odor is represented by an exponential distribution of firing rates, with the same mean for all odors and all odor concentrations. In Step 2, the PNs expand the dimensionality, projecting to 2000 KCs, connected by a sparse, binary random projection matrix. In Step 3, the KCs receive feedback inhibition from the anterior paired lateral (APL) neuron, which leaves only the top 5% of KCs to remain firing spikes for the odor. This 5% corresponds to the tag (hash) for the odor. B) *Illustrative odor responses.* Similar pairs of odors (e.g., methanol and ethanol) are assigned more similar tags than dissimilar odors. Darker shade denotes higher activity. C) *Differences between conventional LSH and the fly’s algorithm.* In the example, the computational complexity for LSH and the fly are the same. The input dimension *d* = 5. LSH computes *m* = 3 random projections, each of which requires 10 operations (5 multiplications + 5 additions). The fly computes *m* = 15 random projections, each of which requires 2 addition operations. Thus, both require 30 total operations.

The second step, where the main algorithmic insight begins, involves a 40-fold expansion in the number of neurons: 50 PNs project to 2000 Kenyon cells (KCs), connected by a sparse, binary random connection matrix (*14*). Each Kenyon cell receives and sums the firing rates from about 6 randomly selected PNs (*14*). The third step involves a winner-take-all circuit using strong inhibitory feedback from a single inhibitory neuron, called APL. As a result, all but the highest firing 5% of Kenyon cells are silenced (*3, 5, 6, 15*). The firing rates of the remaining 5% corresponds to the tag assigned to the input odor.

## Our contributions

From a computer science perspective, we view the fly’s circuit as a *hash function*, whose input is an odor and whose output is a tag (called a *hash*) for that odor. While tags should discriminate odors, it is also to the fly’s advantage to associate very similar odors with similar tags (Figure 1B), so that conditioned responses learned for one odor can be applied when a very similar odor, or a noisy version of the learned odor, is experienced. This led us to conjecture that the fly’s circuit produces tags that are *locality-sensitive*; i.e., the more similar a pair of odors, the more similar their assigned tags. Interestingly, locality-sensitive hash functions (LSH (*16–19*)) serve as the foundation for solving numerous similarity search problems in computer science. The fly’s algorithm, however, departs from traditional LSH algorithms in three ways (discussed later): it uses *sparse, binary* random projections to *expand* the dimensionality of the input, and it then *sparsifies* the tag using a winner-take-all circuit.

Here, we investigate this connection, offering the following contributions:

1. We translate insights from the fly’s circuit to develop a new class of LSH algorithms for efficiently finding approximate nearest-neighbors of high-dimensional points (Figure 1).
2. We prove analytically that the fly’s circuit constructs tags that preserve the neighborhood structure of inputs (as defined under the *ℓ* _2_ norm). The fly’s approach, however, is much more computationally efficient than common approaches often used in the literature.
3. We show empirically that the fly’s algorithm improves performance or computational efficiency of finding nearest neighbors versus traditional LSH algorithms on three benchmark datasets (Figures 2 and 3).

We conclude by describing how the fly’s core procedure to construct odor tags might also illuminate the logic of several vertebrate neural circuits (Table 1).

## The relationship amongst nearest-neighbor search, locality-sensitive hashing, and the fly olfactory circuit

Imagine you are provided an image of an elephant and seek to find the 100 images — out of the billions of images on the Web — that look most similar to your elephant image. This is called the *nearest-neighbors search problem*, which is of fundamental importance in information retrieval, data compression, and machine learning (*17*). Each image is typically represented as a *d*-dimensional feature vector that lies at a point in ℝ^*d*^ space. (Recall that each odor a fly processes is located at a point in 50-dimensional space, ℝ^*50*^_+_). A distance metric is used to compute the similarity between two images (feature vectors), and the goal is to efficiently find the nearest-neighbors of any query image. If the Web contained only a few images, then brute force linear search could easily be used to find the exact nearest neighbors. If the Web contained many images, but each image was represented by a low-dimensional vector (e.g., 10 or 20 features), then space partitioning methods, such as *kd*-trees (*20*), would similarly suffice. However, for large databases with high-dimensional data, neither approach scales (*19*).

Fortunately, in many applications, returning an *approximate* set of nearest-neighbors that are “close enough” to the query is adequate, so long as they can be found quickly. This has motivated an approach for finding approximate nearest-neighbors using a probabilistic technique called locality-sensitive hashing (LSH (*17*)). For the fly, the “tag” (or hash) of an odor corresponds to the vector of Kenyon cell firing rates for that odor. The locality-sensitive property states that two odors that are similar (e.g., methanol and ethanol) will be represented by two tags that are themselves similar (Figure 1B). Likewise, for image search, the tag of an elephant image will be more similar to the tag of another elephant image than to the tag of a skyscraper image. Formally:

**Definition 1.** *A hash function h* : ℝ^*d*^ *→* ℝ^*m*^ *is called locality-sensitive if for any two points p, q ∈* ℝ^*d*^, Pr[*h*(*p*) = *h*(*q*)] = *sim*(*p, q*)*, where sim*(*p, q*) *∈* [0, 1] *is a similarity function defined on two input points.*

Unlike a traditional (non-LSH) hash function, where the input points are scattered randomly and uniformly over the range, the LSH function *h* provides a distance-preserving embedding of points from *d*-dimensional space into *m*-dimensional space (the latter corresponds to the tag). Thus, points that are closer to one another in input space have a higher probability of being assigned the same or similar tag than points that are far apart^1^.

To design a LSH function, one common trick is to compute random projections of the input data (*16–19*) — i.e., to multiply the input feature vector by a random matrix. The Johnson-Lindenstrauss lemma (*16, 21–23*), and its many variants (*24–26*), provide strong theoretical bounds on how well locality is preserved when embedding data from *d* into *m* dimensions using various types of random projections.

Strikingly, the fly also assigns tags to odors by using random projections (step 2, 50 PNs *→* 2000 KCs), which provides a key clue towards the function of this part of the circuit. There are, however, three differences between the fly’s algorithm versus conventional LSH algorithms. First, the fly uses *sparse, binary* random projections, whereas LSH functions typically use dense, i.i.d. Gaussian random projections that are much more expensive to compute. Second, the fly *expands* the dimensionality of the input after projection (*d ≪ m*), whereas LSH contracts the dimension (*d ≫ m*). Third, the fly *sparsifies* the higher-dimensionality representation using a winner-take-all mechanism, whereas LSH preserves a dense representation.

## Deriving the distance-preserving properties of the fly’s olfactory circuit

We can view the mapping from projection neurons (PNs) to Kenyon cells (KCs) as a bipartite connection matrix, with *d* = 50 PNs on the left and the *m* = 2000 KCs on the right. The nodes on the left take values *x*_1_*,…, x*_*d*_ while those on the right are *y*_1_*,…, y*_*m*_. Each value *y*_*j*_ is equal to the sum of a small number of the *x*_*i*_’s; we represent this relationship by an undirected edge connecting every such *x*_*i*_ with *y*_*j*_. This bipartite graph can be summarized by an *m × d* adjacency matrix *M* :

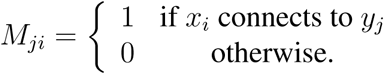

Moving to vector notation, with *x* = (*x*_1_*,…, x*_*d*_) *∈* ℝ^*d*^ and *y* = (*y*_1_*,…, y*_*m*_) *∈* ℝ^*m*^, we have^2^:

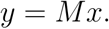

After feedback inhibition from the APL neuron, only the *k* highest firing KCs retain their values; the rest are zeroed out. This winner-take-all mechanism produces a sparse vector *z ∈* ℝ^*m*^ (called the tag) with:

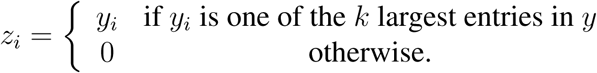

A simple model of *M* is a sparse, binary random matrix: each entry *M*_*ij*_ is set independently with probability *p*. Choosing *p* = 6*/d*, for instance, would mean that each row of *M* has roughly 6 ones (and all of the other entries are 0), which matches experimental findings (*14*).

In the Supplement, we prove that the first two steps of the fly’s circuitry produces tags that preserve *ℓ*_2_ distances of input odors in expectation:

**Lemma 1.** *If two inputs x, x′ ∈* ℝ^*d*^ *get projected to y, y′ ∈* ℝ^*m*^*, respectively, we have*

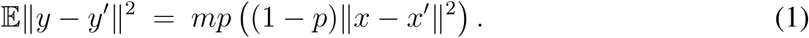

The third step (winner-take-all) is then a simple method for sparsifying the representation while preserving the largest and most discriminative coefficients (*3, 27*).

In the Supplement, we also prove that when *m* is large enough (i.e., the number of random projections is *O*(*d*)), the variance ∥*y*∥^2^ is tightly concentrated around its expected value, such that

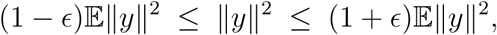

with high probability, for small *ϵ >* 0.

Together, these results prove that the fly’s circuit represents a new LSH family.

## Empirical evaluation on 3 benchmark datasets

To perform a fair comparison between the fly’s algorithm and traditional LSH (*16–19*), we fixed the computational complexity of both algorithms to be the same (Figure 1C). That is, the two approaches were fixed to use the same number of mathematical operations to generate a hash of length *k* (i.e., a vector with *k* non-zero values) for each input. See Materials and Methods for details.

We compared the two algorithms for finding nearest-neighbors on three benchmark datasets: SIFT (*28*) (*d* = 128), GLOVE (*29*) (*d* = 300), and MNIST (*30*) (*d* = 784). SIFT and MNIST both contain vector representations of images used for image similarity search, whereas GLOVE contains vector representations of words used for semantic similarity search. We collected a subset of each dataset with 10,000 inputs each, where each input was represented as a feature vector in ℝ^*d*^. To test performance, we selected 1000 random query inputs from the 10,000 and compared true versus predicted nearest-neighbors. That is, for each query, we found the top 2% (200) of its true nearest-neighbors in input space, determined based on the Euclidean distance between feature vectors. We then found the top 2% of predicted nearest-neighbors in hash space (i.e., the range of *h*), determined based on Euclidean distance between tags (hashes). We varied the length of the hash (*k*) and computed the overlap between the ranked lists of true and predicted nearest-neighbors using the mean average precision (*31, 32*). We averaged the mean average precision over 50 trials, where in each trial the random projection matrices and the queries changed.

Below, we isolated each of the three differences between the fly’s algorithm and LSH to test their individual effect on nearest-neighbors retrieval performance.

## Sparse binary vs. dense Gaussian random projections (Figure 2A)

**Figure 2:**
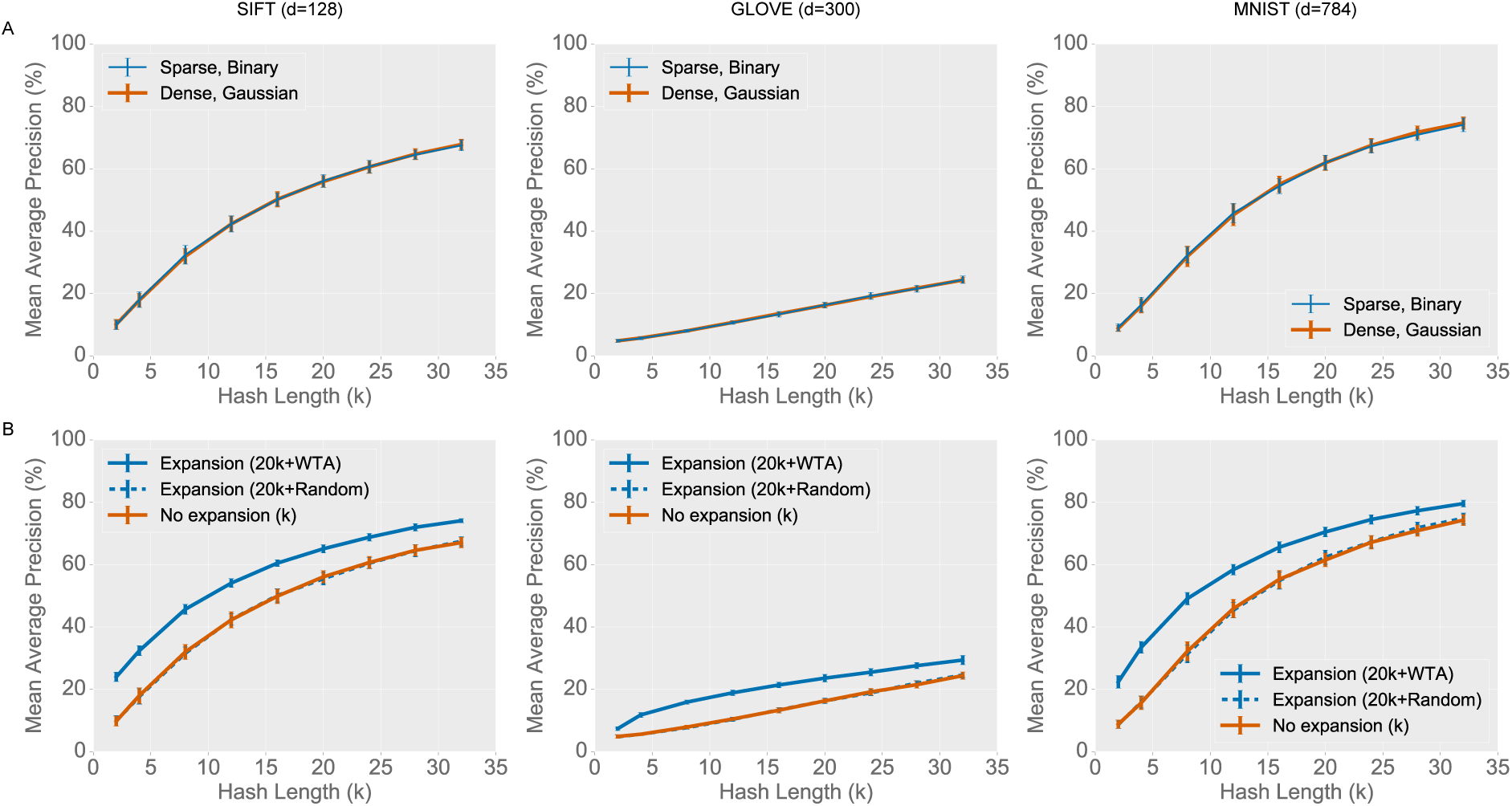
Empirical comparison of different random projection types and tag-selection methods. In all plots, the *x*-axis is the length of the hash, and the *y*-axis is the mean average precision (higher is better). A) Sparse, binary random projections offer near-identical performance as dense, Gaussian random projections, but the former provide a large savings in computation. B) The expanded-dimension (from *k →* 20*k*) plus winner-take-all sparsification further boosts performance versus non-expansion. Results are consistent across all three benchmark datasets. Error bars indicate standard deviation over 50 trials.

First, we simply modified LSH to use sparse binary, instead of dense Gaussian, random projections. No other aspect of the fly’s algorithm was used. We found that the two random projection types produced nearly identical retrieval performance on all three datasets and across all hash lengths (Figure 2A). These results support our theoretical calculations that the fly’s random projection is indeed locality-sensitive. Moreover, the sparse, binary random projection achieved a computational savings of a factor of 20 versus the dense, Gaussian random projection (SI Text).

We also varied the number of input indices (PNs) each Kenyon cell samples, from 1%, to 10% (6 out of 50, as the fly does), to 50%. We found the most consistent performance when sampling 10%, with no improvement in performance at 50% (Figure S1).

## Winner-take-all (WTA) vs. random tag selection following the expansion (Figure 2B)

Sec-ond, we implemented the full fly’s algorithm and compared different methods to select Kenyon cells (KCs) that constitute the tag. Here, we used 20*k* random projections for the fly to equate the number of mathematical operations used by both algorithms (SI Text). We found much better performance using WTA, which selects the top *k* firing KCs as the tag, versus a selection of *k* random KCs (Figure 2). For example, on the SIFT dataset with hash length *k* = 4, random selection yielded a 17.7% mean average precision versus roughly double that (32.4%) using the WTA. Thus, selecting the top firing neurons best preserves relative distances between inputs; the increased dimension also makes it easier to segregate dissimilar inputs. For random tag selection we selected *k* random (but fixed for all inputs) KCs for the tag; hence, its performance is effectively the same as just doing *k* random projections, as in LSH.

## Overall comparison between the fly and LSH (Figure 3)

**Figure 3:**
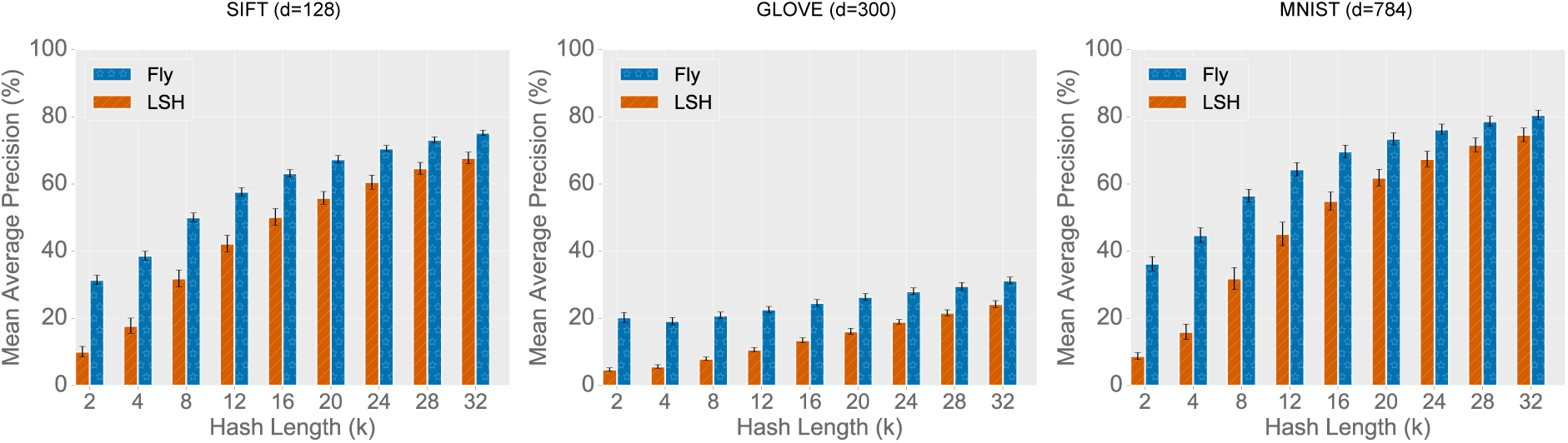
Overall comparison between the fly’s algorithm and locality-sensitive hashing (LSH). In all plots, the *x*-axis is the length of the hash, and the *y*-axis is the mean average precision (higher is better). A 10*d* expansion was used for the fly. Across all three datasets, the fly’s method outperforms LSH, most prominently for short codes. Error bars indicate standard deviation over 50 trials.

Third, to more closely mimic the fly’s circuitry, we implemented the full fly’s algorithm but with a further expansion of the dimensionality from 20*k* to 10*d* Kenyon cells. Overall, we found significant gains compared to LSH across all three datasets (Figure 3). The gains were highest for very short codes, where we see an almost 3x improvement in mean average precision (e.g., for MNIST with *k* = 4, 16.0% for LSH versus 44.8% for the fly).

We also used the fly’s algorithm to implement binary locality-sensitive hashing (*33, 34*), where the LSH function *h* : ℝ^*d*^ *→* ℤ^*m*^. In other words, instead of using the *values* of the top *k* Kenyon cells as the tag, we used their *indices*. The fly’s method improved over a common prior approach, which binarizes each Kenyon cell to 0 or 1 based on whether its value is *≤* 0 or *>* 0, respectively (Figure S3). This suggests that the fly’s ingredients may also be useful across other LSH families.

Finally, the fly’s algorithm is also scalable. While designed biologically for *d* = 50, our datasets included dimensionality up to *d* = 784 (MNIST). We also tested the fly’s algorithm on the GIST image dataset (*28*), where *d* = 960, and found a similar trend in performance (Figure S2).

## Discussion

Overall, we identified a new brain algorithm that supports an important sensory function (olfaction); we derived its distance-preserving properties theoretically; and we empirically evaluated its performance for finding nearest-neighbors on several benchmark datasets. Our work offers a new synergy between strategies for similarity matching in the brain (*35*) and algorithms for nearest-neighbor search in large-scale information retrieval systems. Our work may also have applications in duplicate detection, clustering, and energy-efficient deep learning (*36*).

There are numerous extensions to LSH (*37*), including using multiple hash tables (*19*) to boost precision (we used just one here for both algorithms), using multi-probe (*38*) so that similar tags can be grouped together into the same bin (which may be easier to implement for the fly since tags are sparse), and various quantization tricks for discretizing hashes (*39*). There are also methods to speed up the random projection multiplication — both for LSH, using fast Johnson-Lindenstrauss transforms (*40, 41*), and for the fly, using fast sparse matrix multiplication. Our goal here was to fairly compare two conceptually different approaches for the nearest-neighbors search problem; in practical applications, all of these extensions will need to be ported to the fly’s algorithm.

Following the fly, we focused on *data-independent* hashing; i.e., hash functions that do not learn from prior data nor use prior data in any way when deriving the tag. Recently, many classes of *data-dependent* LSH families have been proposed, including PCA hashing (*42*), spectral hashing (*43*), semantic hashing (*44*), deep hashing (*32*), and others (*45*) (reviewed by Wang et al (*46*)). Some of the fly’s ingredients have been used piece-meal before; for example, MinHash (*47*) and winner-take-all hash (*48*) both use WTA-like components, though neither propose expanding the dimensionality; similarly, random projections are used in many LSH families, but none, to our knowledge, use sparse, binary projections. The combination of these computational ingredients thus appears novel, and it seems remarkable to us that evolution has discovered them for fly olfaction.

Finally, while the fly olfactory system has been extensively mapped experimentally, there is some evidence that the three hallmarks used in the fly’s circuit motif may also appear in other brain regions and species (Table 1). Thus, locality-sensitive hashing may be a general principle of computation used in the brain (*49*).

**Table 1:** The generality of locality-sensitive hashing in the brain.

|  | **Step 1** | **Random Proj.** | **Step 2 (Expansion)** | **Step 3 (WTA)** |
| --- | --- | --- | --- | --- |
| Fly olfaction | Antennae lobe<br>50 glomeruli | Sparse, binary<br>Samples 6 | Mushroom body<br>2000 Kenyon cells | APL neuron<br>top 5% |
| Mouse olfaction | Olfactory bulb<br>1000 glomeruli | Dense, weak<br>Samples all | Piriform cortex<br>100K semi-lunar cells | Layer 2A<br>top 10% |
| Rat cerebellum | Pre-cerebellar<br>nuclei | Sparse, binary<br>Samples 4 | Granule cell layer<br>100M granule cells | Golgi cells<br>top 10–20% |
| Rat hippocampus | Entorhinal cortex<br>30K grid cells | Unknown | Dentate gyrus<br>1.2M granule cells | Hilar cells<br>top 2% |

The steps used in the fly olfactory circuit and their potential analogs in vertebrate brain regions.

## Footnotes

1 In practice, a second (traditional) hash function is used to place each *m*-dimensional point into a discrete bin so that all similar images lie in the same bin, for easy retrieval. In this paper, we focus only on designs for the LSH function, *h*.

2 In practice, an additional quantization step is used for discretization: *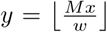*, where *w* is a constant, and ⌊*·* ⌋ is the floor operation.

